# Quantification and potential functional relevance of binding cooperativity of adjacent transcription factors on DNA

**DOI:** 10.1101/2024.11.20.624593

**Authors:** Xinyao Wang, Chen Xie, Ke Shen, Dubai Li, Xiaoliang Sunney Xie

## Abstract

In eukaryotes, expression of a particular gene is regulated by a combination of transcription factors (TFs) bound on regulatory regions of the genomic DNA (promoters and enhancers). Recent advances in genomic sequencing technology have allowed measurements of TFs’ footprints and binding affinities on DNA at the single-molecule level, permitting the probing of binding cooperativity among adjacent TFs. This necessitates quantitative descriptions of TFs’ binding cooperativity and understanding of its potential functional relevance. In this study, we show that, thermodynamically, the binding affinities of two adjacent TFs can either increase together (positive cooperativity) or decrease together (negative cooperativity), rather than changing in opposite directions. Their binding cooperativity can be quantified by the *γ* coefficient, which is independent of TF concentrations, and can be determined by single-molecule binding reads either from *in vitro* thermodynamic condition or cellular condition of non-equilibrium steady state (NESS). The functional relevance of positive cooperativity, which has been extensively discussed in the literature, is the sigmoidal binding curve around a TF concentration threshold (analogous to oxygen binding to hemoglobin), whereas the functional relevance of negative cooperativity is two-fold. First, mutual exclusion of the two TFs enables bidirectional gene switching similar to CI-Cro system in phage *λ*. Second, under a non-equilibrium steady-state condition, in which TFs often exhibit intranuclear concentration fluctuations, negative binding cooperativity assures fast TF dissociation from DNA and hence rapid response for gene expression regulation.

**Significance Statement:** In eukaryotes, multiple transcription factors (TFs) bind to regulatory regions of DNA to control gene expression. The binding affinity of one TF to a specific DNA site is influenced by the binding of another TF at an adjacent site, exhibiting either positive or negative cooperativity (the TFs either strengthen or weaken each other’s binding). Here we present metric for such cooperativity from experimental measurables. Functionally, positive cooperativity assures sensitive response above a threshold of TF concentrations, whereas functional roles for negative cooperativity might be two folds: First, mutual exclusion of TFs’ binding enables bidirectional gene switching. Second, as TF concentrations oscillate under non-equilibrium steady-state condition, negative binding cooperativity assures fast TF dissociation, hence rapid switching of transcription.

---

Binding cooperativity among biomolecules is fundamental to many biological processes, and the classic example is the binding of multiple oxygen molecules to hemoglobin (1). Binding cooperativity can be either positive and negative, the former is exemplified by hemoglobin’s oxygen binding (1), and the latter has received increased attention (2–4), though its functional relevance remains elusive. Multiple TFs bind to DNA regulatory regions, such as promoters and enhancers, to control gene expression (5–7). Mechanistically, positive TF binding cooperativity arises from protein-protein interactions among TFs (8–10), which recruits multiple proteins to assemble functional complexes. Negative TF binding cooperativity may result from competitive binding of TFs to closely-spaced binding sites on DNA (11). Furthermore, allostery through DNA can lead to either positive or negative binding cooperativity, which is dependent on the distance between the TFs, and varies with a periodicity of ∼ 10 base pairs (12, 13).

Traditionally, TF binding cooperativity can be detected by biochemical bulk measurements such as electrophoretic mobility shift assay (EMSA) (14, 15). Recently, single-molecule imaging (12, 13) and high-throughput sequencing techniques such as CAP-SELEX (16), single molecule footprinting (SMF) (17), Fiber-seq (18), and footprinting with deaminase (FOODIE) (7) offer the promise of detecting TF binding cooperativity with improved accuracy. A key challenge thus arises: How do we quantify TF binding cooperativity from the single-molecule data from the above techniques? Moreover, what are the functional relevance of positive and negative cooperativity? This paper addresses these critical questions.

## Results

Sequencing techniques that detect transcription factor (TF) binding at the single-molecule level are typically applied to groups of identical cells (*in vivo* experiments) or to purified DNA molecules with one or a few TFs (*in vitro* experiments). In both cases, the initial step involves identifying TF binding sites at the ensemble level.

A footprint refers to a short DNA sequence bound by a TF, characterized by a specific binding fraction. Subsequently, the cooperative binding between adjacent footprints (i.e., TF binding sites) can be statistically or thermodynamically quantified. For any two adjacent TF binding sites, the numbers of DNA molecules that are co-bound (*a*), singly bound (either *b* or *c*), and unbound at both sites (*d*) are recorded (Fig 1A). In the following section, we will define and calculate several statistical and thermodynamic metrics, then discuss their relationship.

**Fig. 1.**
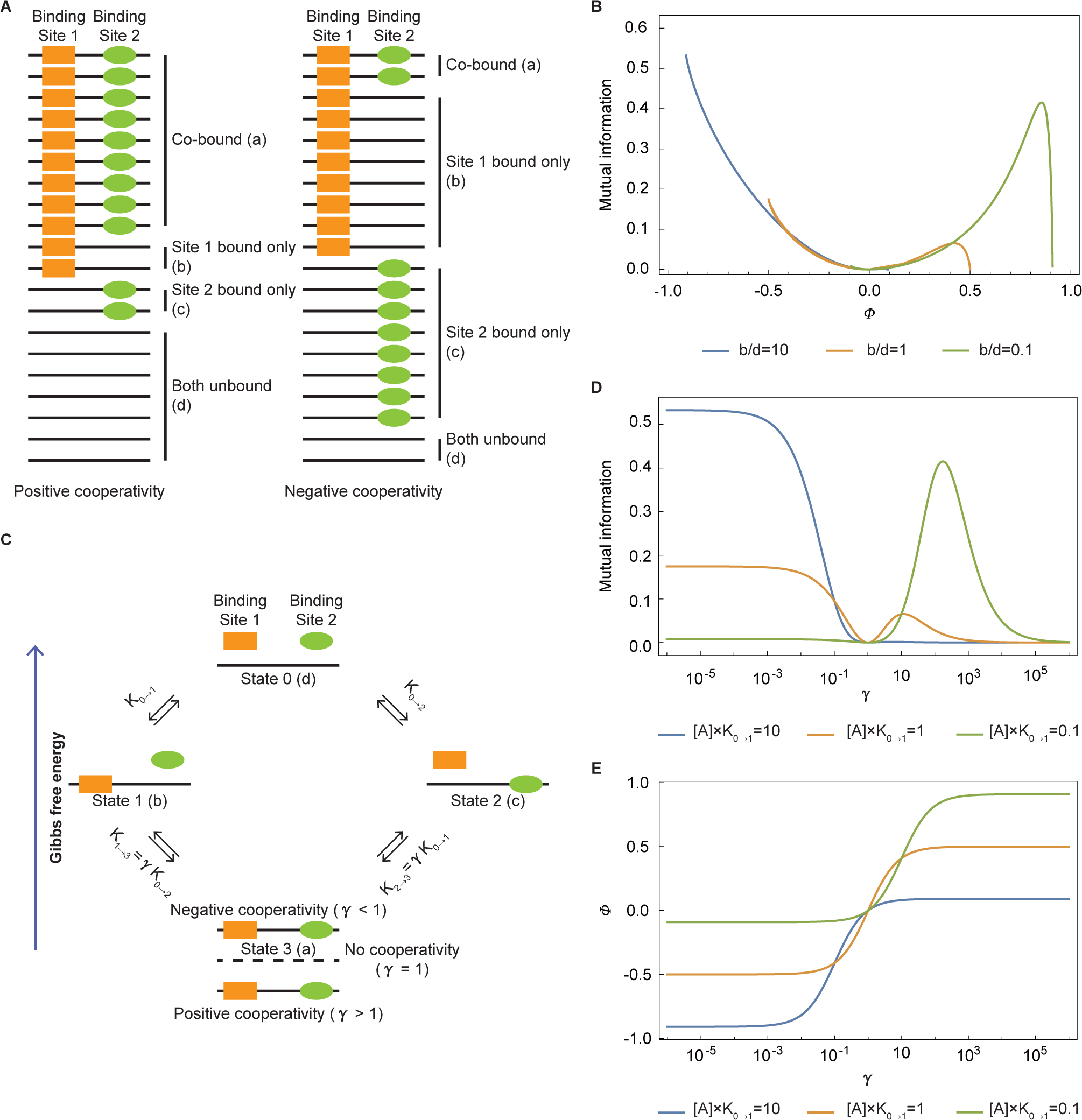
Relationship between *Φ*, mutual information (*MI*) and *γ*. (A) Schematics of single molecules with two adjacent binding sites, showing scenarios of positive cooperativity (left) and negative cooperativity (right). *a* to *d* are single-molecule counts. (B) The relevance between *MI* and *Φ* for a special case of Fig 1A where identical TFs bind at two identical sites (i.e., *b* = *c*). All three situations are shown: the binding fraction 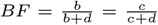 of one site given the adjacent site not bound is high (blue), medium (orange), and low (green). (C) Thermodynamic cycle illustrating the binding of two TFs at adjacent DNA sites. Four possible DNA binding states (0–3) are depicted, with single-molecule counts shown in parentheses. Binding constants are indicated, where positive cooperativity enhances the association equilibrium constant (*γ >* 1) and negative cooperativity reduces it (*γ <* 1). Orange rectangle: TF A; green oval: TF B; Black straight line: DNA. (D) The relevance between *MI* and *γ* for the special case of Fig 1C where identical TFs bind on two identical sites. Binding fractions 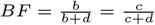of one site given the adjacent site not bound are presented under high (blue), medium (orange), and low (green) conditions. (E) Relationship between *Φ* and *γ* corresponding to the case in Fig 1C with identical TFs bind on two identical sites, showing three different binding fraction 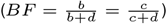 conditions (high, medium, and low, indicated by blue, orange, and green respectively).

### *Φ* Correlation Coefficient

From a statistical perspective, two binding sites can be modeled as two binary random variables:

TF Binding Site 1 and TF Binding Site 2, each of which has two possible states: bound and unbound. The *Φ* coefficient (equation 1) is commonly used to quantify the correlation between these two variables, which is due to the cooperativity between the two TF binding sites.

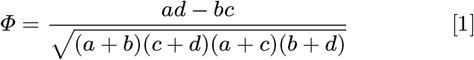

The *Φ* coefficient is analogous to the Pearson correlation coefficient in its statistical interpretation. Through simple derivation, we obtain:

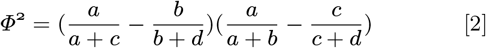

The four fractions in equation 2 represent the binding fractions of Site 1 given Site 2 bound and unbound, and the binding fractions of Site 2 given Site 1 bound and unbound, respectively. Consequently, *Φ* quantifies the overall TF binding fractions’ correlation between the two sites and has concentration dependence. *Φ* ranges from -1 to 1, it can be interpreted as 0 for no cooperativity, 1 for completely positive cooperativity (“all or none”), -1 for completely negative positivity (“either or”), and nominally between 0 and 1 for partially positive cooperativity and between -1 and 0 for partially negative cooperativity.

In the analysis of real data, the availability of total DNA molecules is often limited. Thus, it is essential to determine whether *Φ* is statistically significantly above or below zero, indicating significant positive or negative cooperativity. Fisher’s exact test of independence is the preferred method for this analysis. If its p-value is below a specified cutoff (such as the commonly used 0.05), then TF binding cooperativity is considered statistically significant. Bootstrapping analysis can further generate a distribution of *Φ*, and if zero is not inside a certain confidence interval (such as the commonly used 95%), then TF binding cooperativity is significant.

### Mutual Information (*MI*)

The statistical significance of *Φ* and TF binding cooperativity can be further understood theoretically through mutual information (*MI*, equation 3) from information theory (19). Here, *MI* quantifies the binding status gained at one binding site by observing binding status at the other site. By definition, *MI* is non-negative. When *MI* is zero, it indicates no information, due to either no cooperativity, or the estimation error of *Φ* is as large as infinite, i.e., impossible to make any conclusion on TF binding cooperativity no matter how many DNA molecules are measured. A small *MI* implies limited information, requiring a large number of DNA molecules to detect significant cooperativity. Conversely, a high *MI* indicates substantial information, allowing for the detection of significant cooperativity with a relatively smaller number of DNA molecules. For a fixed number of DNA molecules, higher *MI* values correspond to lower p-values in Fisher’s exact test.

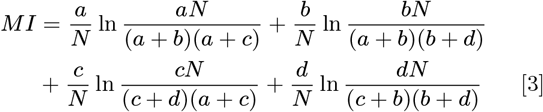

The relationship between *MI* and *Φ* is further illustrated in Fig 1B, where it is discussed in a simplified scenario that identical TFs bind on two identical sites (i.e., *b* = *c*). Three conditions are shown: a high binding fraction 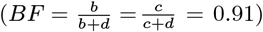 with *b/d* = 10 (blue line), a medium binding fraction with *BF* = 0.5 (*b/d* = 1, orange line), and a low binding fraction with *BF* = 0.09 (*b/d* = 0.1, green line). Under all conditions, *MI* generally increases as |*Φ*| increases for most *Φ* values, consistent with an enhanced ability to detect larger |*Φ*| values. However, at the extreme positive values of *Φ, MI* becomes very small. This occurs because the power to detect significantly positive *Φ* is low when nearly all DNA molecules are co-bound (with *b, c*, and *d* approaching zero).

### Thermodynamics of TF Binding Cooperativity and *γ* Coefficient

In the preceding discussion, we examined the quantification of TF binding cooperativity using the *Φ* coefficient and *MI* from statistics and information theory, but *Φ* is concentration dependent and lack of thermodynamical interpretation. To address this limitation, we now present a thermodynamic model describing the binding of two TFs at adjacent sites on DNA (Fig 1C). For each reversible reaction, we define *k*_+_ and *k*_−_ as the kinetic rate constants for the forward binding and reverse dissociation reactions, respectively. When this system reaches thermodynamic equilibrium, the forward binding reaction is balanced by the reverse dissociation reaction, and the binding affinity represented by the association binding constant 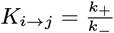(*i, j* stand for different states) is defined as follows ([*A*] or [*B*] represents the concentration of TF A or TF B):

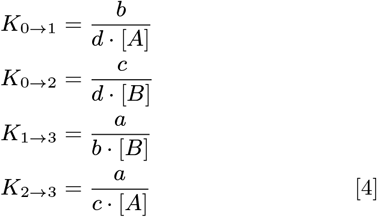

Given that Gibbs free energy is a state function and thus independent of the reaction pathway, we have:

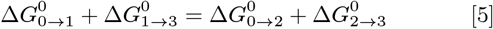

Where 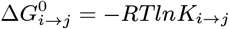, thus we obtain:

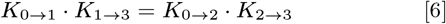

This thermodynamic cycle has also been previously discussed (12). Here, we further introduce *γ* to quantify these circumstances:

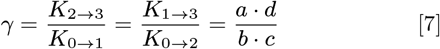

By definition, the *γ* coefficient is independent of TF concentrations and ranges from 0 to positive infinity, with values indicating the degree of cooperativity. *γ* = 1 indicates that the binding constant of TF B in the presence of TF A is equal to its binding constant in the absence of TF A, signifying independent binding of TF A and TF B on DNA. When *γ >* 1, this declares the binding affinities of TF A and TF B increase together (positive cooperativity). The existence of positive cooperativity causes relatively lower Gibbs free energy (more stable, Fig 1C).

The positive cooperativity can be caused by TF complexes constituted by protein-protein interactions (8–10). conversely, when *γ <* 1, the binding affinities of TF A and TF B decrease together (negative cooperativity). Negative cooperativity results in relatively higher Gibbs free energy (less stable, Fig 1C). The *γ* coefficient is commonly known as the odds ratio in statistics. Odds ratio (20) quantifies the strength of the relationship between two conditions (binding events of Site 1 given Site 2 bound, and unbound).

The negative cooperativity can arise from competitive binding among TFs with closely adjacent binding sites (11). Moreover, the cooperativity between two TFs binding at adjacent sites can oscillate between positive and negative situations with a period of 10 base pairs due to DNA allostery (12, 13). Compared to the widely used Hill coefficient, *γ* provides a more intuitive and perceivable assessment for binding cooperativity between two TFs. For convenience, log *γ* can also be used to quantify TF binding cooperativity, with zero corresponding no cooperativity, log *γ >* 0 for positive cooperativity and log *γ <* 0 for negative cooperativity.

The previous discussion about *γ* focuses on the case of thermodynamic equilibrium, which is usually satisfied in the *in vitro* experiments. However, cells under non-equilibrium steady-state (NESS) condition (21) often exhibit TF concentration fluctuation (22, 23), the timescale of which is rather long, on the order of an hour. On the other hand, the dynamic binding and unbinding of TFs on the DNA takes place on the much shorter timescale, from seconds to minutes (24). This separation of timescales, combined with the fact that *γ* is independent of TF concentrations, implies that the measurement of *γ* from single-molecule reads (a, b, c, and d) of genomic DNAs should reflect the intrinsic thermodynamic property of *γ*, and is independent of TF concentration fluctuation under the NESS condition of cells.

### Relationships among *Φ, MI*, and *γ*

In the analysis of real data, as discussed for the *Φ* coefficient, bootstrapping analysis for *γ* can be performed in the same way. If one is not inside a certain confidence interval (such as the commonly used 95%), then TF binding cooperativity is significant.

The statistical significance of *γ* can also be theoretically evaluated using *MI*. As illustrated in Fig 1D, we simplify the scenario to two identical TFs binding at two identical sites, then we consider three binding conditions: the binding fraction 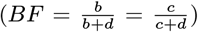 of one site given the adjacent site not bound is high ([*A*]*K*_0→1_ = *b/d* = 10, *BF* = 0.91), medium ([*A*]*K*_0→1_ = *b/d* = 1, *BF* = 0.5), and low ([*A*]*K*_0→1_ = *b/d* = 0.1, *BF* = 0.09). The patterns of *γ* in these scenarios become more complicated. The expectation is that the power to significantly detect larger |ln *γ*| is larger, i.e., *MI* becomes larger when |ln *γ*| becomes larger. However, more situations violate this expectation. For example, under high binding fraction conditions, *MI* remains near zero for *γ >* 1. In the medium and low binding fraction cases, *MI* becomes very small when *γ* becomes very large. These are due to that the power to significantly detect positive ln *γ* is very small when almost all the DNA molecules are co-bound (b, c, and d are zero or almost zero). Conversely, under low binding fraction conditions, *MI* for *γ <* 1 is always almost zero. This is due to that the power to significantly detect negative ln *γ* is very small when almost all the DNA molecules are unbound at both sites (a, b, and c are zero or almost zero).

So far, we have introduced *Φ* and *γ* to quantify cooperativity, and we now examine their relationship. With a simple derivation, dividing both the numerator and denominator in equation 1 by *bc* and substituting *a, b, c*, and *d* with TF concentrations and binding constants (see Extended Methods), *Φ* coefficient can be expressed in terms of *γ*, TF concentrations and binding constants as equation 8.

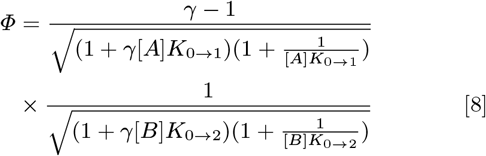

It is evident that the *Φ* coefficient is influenced not only by variations in binding affinity (*γ*) but also by TF concentrations and binding constants. This relationship is further illustrated in Fig 1E with the same simplification and conditions in Fig 1D. In all scenarios, *Φ* increases monotonically with *γ*, and reaches saturation at very high or low values of *γ*. Therefore, *γ* serves as an effective measure of variations in TF binding affinity due to cooperativity, while *Φ* is well-suited for capturing TF binding fraction variations due to cooperativity.

### Functional Relevance of Positive Cooperativity

We first study the functional relevance of positive cooperative binding under thermodynamic equilibrium. To simplify the system, we assume that the two TFs and their binding sites in Fig 1C are identical. The simulation is carried out under these parameters: *k*_+,0→1_=5 *×* 10^5^ M^−1^*·*s^−1^, *k*_−,1→0_=0.02 s^−1^, and 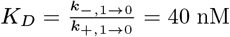, which are close to the TF binding in mammalian cells and consistent with literature (25–27). Fig 2A illustrates the response of the co-binding fraction (State 3 in Fig 1C) to TF concentration with different *γ* values. All response curves follow a sigmoidal pattern, in which co-binding with positive cooperativity (*γ >* 1) exhibiting a more rapid response to changes in concentration, thereby acting as a sensitive response around a threshold of TF concentration.

**Fig. 2.**
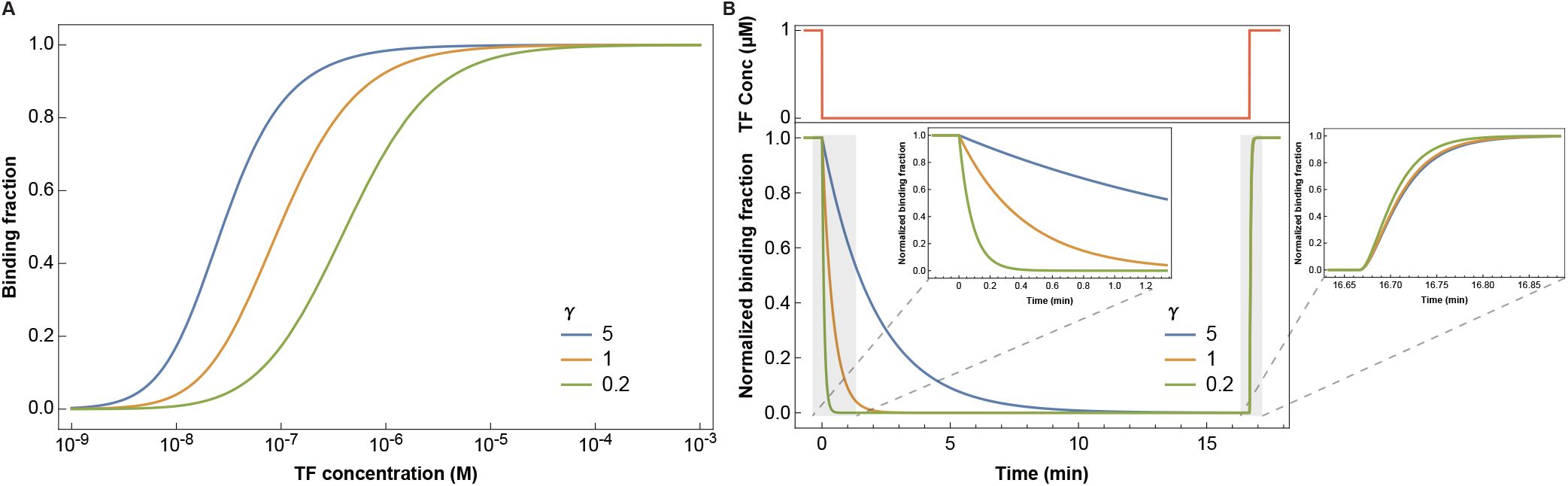
Thermodynamics and kinetics of identical TFs binding at two adjacent DNA sites with identical binding constants (Special case of Fig 1C). (A) TF concentration exhibits sigmoidal dependence of co-binding fraction (Fig 1B, State 3) for different *γ* values under thermodynamic equilibrium states. (B) Response of co-binding fraction to oscillating TF concentration in the form of a rectangular wave. Top panel: TF concentration oscillates between 1μM and 0 as a rectangular wave with a duty cycle of 50% and a period of 1000 s. Left inset: Enlarged view showing co-binding fraction with each of different *γ* values descends with different rates when TF concentration steps down from 1μM to 0. Right inset: Enlarged view showing co-binding fraction with each of different *γ* values ascends with different rate as TF concentration shifts from 0 to 1μM.

### Functional Relevance of Negative Cooperativity

Negative cooperativity implies that the binding of TF A would weaken the binding of TF B, resulting in an either-or tendency for TF A and B to bind DNA. Consequently, negative cooperativity ensures mutual exclusivity in TF binding, akin to the CI-CRO system in phage *λ* (6), and facilitates bidirectional gene switching.

In living organisms, TF concentrations often oscillate in respond to specific cellular and environmental stimuli (22, 28). To explore the functional significance of negative cooperative binding under fluctuating TF concentrations, we commence with a simplified model of Fig 1C which takes *A* and *B* as the same transcription factor and assume two identical binding sites. These assumptions signify that the two binding sites share the same binding affinity 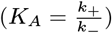, association rate constant (*k*_+_), and dissociation rate constant (*k*_−_). In this scenario, we could derive three time-dependent differential equations with specific boundary conditions (see Extended Methods for details).

To obtain a concise physicochemical model, we introduce several additional assumptions. First, we simply take the change of TF concentration as a rectangular wave function. Second, although both the association rate constant (*k*_+_) and dissociation rate constant (*k*_−_) may vary with cooperativity, we neglect changes in *k*_+_, as it is primarily governed by bimolecular diffusion (29). Thus, we attribute changes in association binding constant entirely to alterations in *k*_−_.

Response of TF co-binding fraction (Fig 1C, State 3) to TF concentration oscillation is simulated under these kinetic parameters: *k*_+,0→1_=5*×*10^5^ M^−1^*·*s^−1^, *k*_−,1→0_=0.02 s^−1^, [*A*] is treated as a rectangular wave oscillating between 1 μM and 0 with a duty cycle of 50% and a period of 1000 s, total DNA concentration is normalized to 1.

Fig 2B illustrates the response of TF co-binding fraction to oscillating TF concentration in the form of a rectangular wave. The simulation starts from thermodynamic equilibrium binding states (binding fraction of different *γ* values are normalized to 1). For *γ* = 0.2 (negative cooperativity), the binding fraction declines rapidly, reaching bottom within 40 s (left inset of Fig 2B), suggesting a fast response to TF concentration fluctuation. In contrast, when *γ* = 5, the co-binding fraction for positive cooperativity reaches bottom much slower than that for negative cooperativity. The right inset of Fig 2B provides a zoom-in view of the response of co-binding fraction when the TF concentration rising from 0 to 1 μM as a step function, due to rapid biomolecular binding. In summary, negative binding cooperativity enables either mutual exclusion of TFs similar to the CI-Cro system in phage *λ* (6) or fast TF dissociation from the DNA in response to a rapid intranuclear TF concentrations change under non-equilibrium steady state conditions, hence a rapid response of transcription.

### Cooperative Binding of Three Transcription Factors (TFs)

In eukaryotic cells, multiple TFs bind to adjacent sites within DNA regulatory regions to regulate gene expression (7, 10). The analyses above merely focus on two TFs’ co-binding on DNA and its statistical quantification, thermodynamics and kinetics. In this section, we extend the discussion to the case of three TFs’ binding at adjacent DNA sites and investigate whether the conclusions drawn from two TFs’ co-binding are applicable to three TFs’ co-binding. Due to the third TF’s binding introduced into the system, the number of possible binding states increases to 2^3^ = 8, making the system way more complicated. The thermodynamic cycles of three TFs’ binding are illustrated in Fig 3. To simplify the analysis, we bring in a new assumption that the binding events of TF C and TF B are independent from each other, as they are not adjacent to each other. This simplifies the system to two distinct cooperativity parameters, *γ*_*CA*_ and *γ*_*AB*_. Under thermodynamic laws, the relationship between the binding constants can be expressed as follows (with a detailed derivation provided in the Extended Methods):

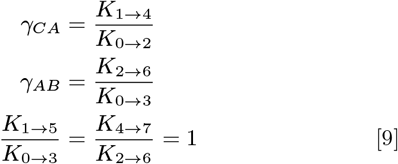

**Fig. 3.**
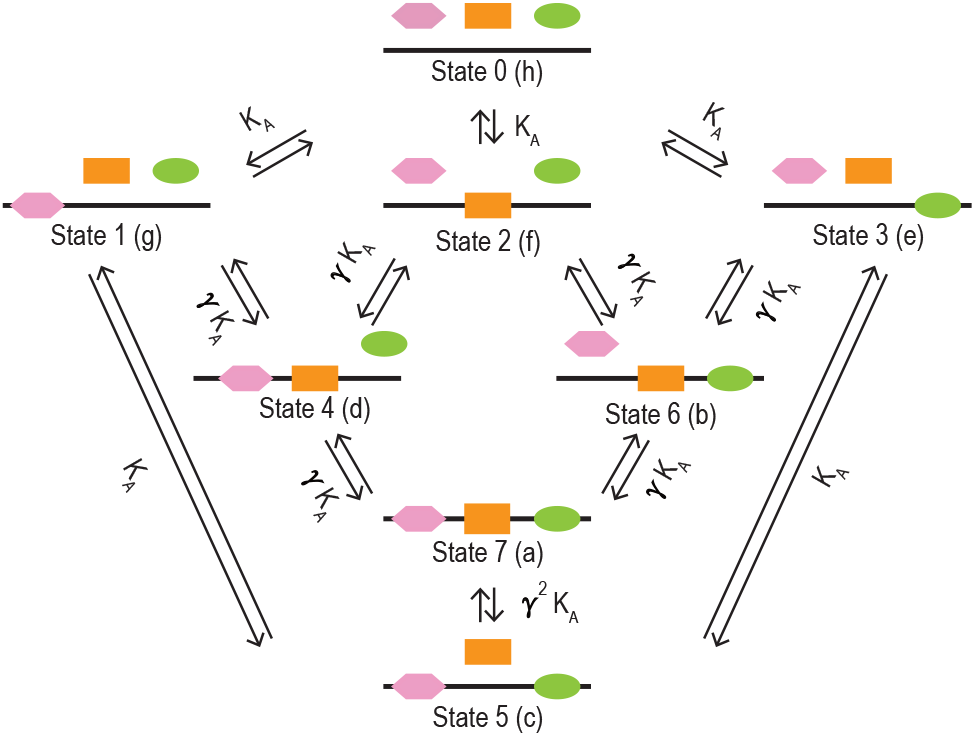
Schematics of thermodynamic cycles of three TFs (C, A, and B) binding at adjacent sites on DNA. Eight DNA binding states (0 - 7) are presented (corresponding single molecule counts are shown in parentheses, which are used for the thermodynamics and kinetics derivation in Extended Methods). Association constants are marked. Orange rectangle: TF A; green oval: TF B; Pink hexagon: TF C; Black straight line: DNA.

Therefore we have:

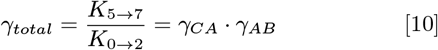

If we set *γ*_*CA*_ = *γ*_*AB*_ = *γ*, then we have *γ*_*total*_ = *γ*^2^. We performed the numerical simulation with this restriction and used the same assumptions and kinetics parameters as those for two TF’s co-binding. Fig 4A shows the steeper sigmoidal concentration dependence of the three TFs’ binding fraction (Fig 3, State 7) than the two TFs’. The binding fraction versus time in response to a rectangular wave function of TF concentration, with different *γ* values are shown in Fig 4B with an analogous conclusion to that of Fig Fig 2B, with an even faster response time from the equilibrium value than two TFs’ case due to third TF’s binding.

**Fig. 4.**
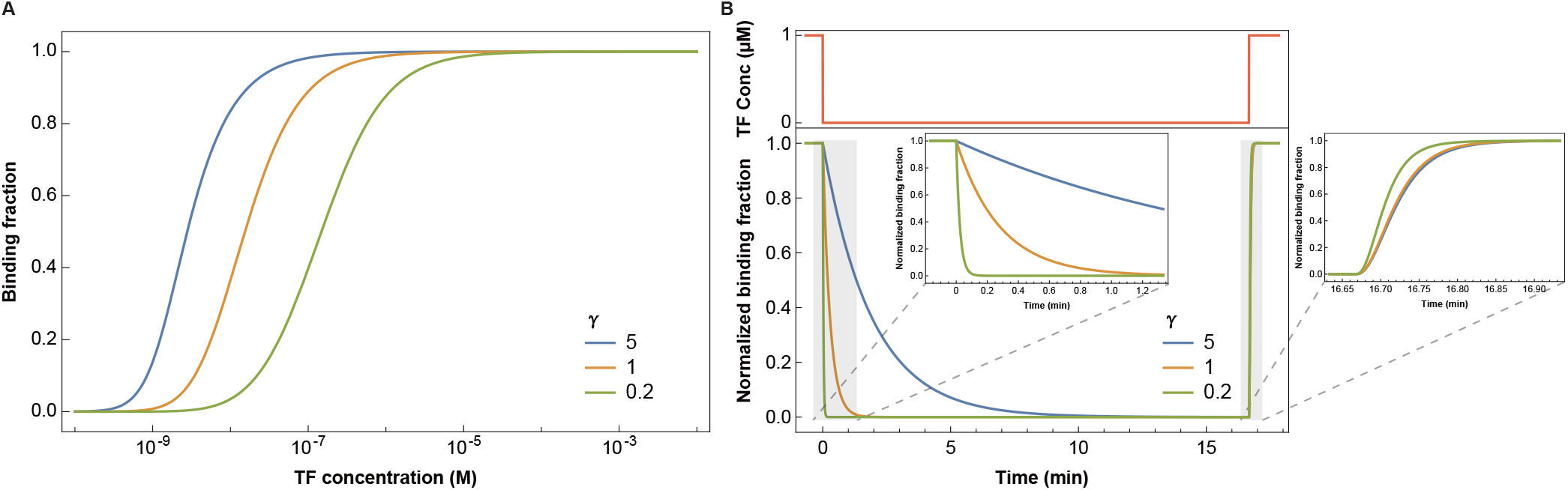
Thermodynamics and kinetics of same TFs binding on three adjacent sites on DNA molecules with identical binding constants (Special case of Fig 3). (A) Dependence of the co-binding fraction (Fig 3, State 7) on TF concentration, exhibiting sigmoidal response for various *γ* values. (B) Response of co-binding fraction to oscillating TF concentration in the form of a rectangular wave. Top panel: TF concentration oscillates between 1μM and 0 as a rectangular wave with a duty cycle of 50% and a period of 1000 s. Left inset: Enlarged view showing co-binding fraction with each of different *γ* values descends with different rates when TF concentration steps down from 1μM to 0. Right inset: Enlarged view showing co-binding fraction with each of different *γ* values ascends with different rate as TF concentration shifts from 0 to 1μM.

## Conclusions

We have introduced three metrics to quantify the TFs’ binding cooperativity: *Φ* coefficient, mutual information (*MI*), and *γ* coefficient. Among these metrics, the *Φ* coefficient captures the correlation between overall binding fractions, but is dependent on TF concentrations. *MI* quantifies how the binding of one site can provide information on the other site, though it does not reflect the strength of cooperativity. The *γ* coefficient is independent of TF concentrations and can serve as an intrinsic metric for TF binding cooperativity. Therefore, we suggest that *γ* is the optimal metric for quantifying TF binding cooperativity. Thermodynamic analysis of *γ* demonstrates that, the binding affinities of the two TFs at adjacent sites either increase together due to positive cooperativity or decrease together due to negative cooperativity, rather than changing in opposite directions. Our thermodynamic investigation further reveals that the functional significance of positive cooperativity lies in its ability to produce sigmoidal TF concentration dependence of the co-binding fraction, thereby enhancing sensitivity. This behavior mirrors the classic receptor-ligand positive cooperativity observed in the hemoglobin-oxygen interaction (1). Given that TF concentrations ubiquitously oscillate in the cell nucleus (22, 28) under non-equilibrium conditions, our kinetics analysis highlights the functional significance of negative cooperativity in assuring rapid TF dissociation from DNA in response to abrupt changes in TF concentration, hence rapid switching of transcription.

## Materials and Methods

The relationship between *Φ* and *γ* is derived in Extended Methods. We derived the thermodynamic cycles of two and three TFs binding at adjacent sites on DNA, with the former shown in the Results section, and the latter is shown in the Extended Methods. The differential equations of the kinetics of two and three TFs binding to adjacent sites on DNA are shown in the Extended Methods.

Mathematica (version 13.0) is used for deriving analytical and numerical solutions and plotting.

## Supporting Information Appendix (SI)

Extended Methods (PDF).

## ACKNOWLEDGMENTS

We thank Wenyang Dong, Runsheng He, Zhi Wang, Zhen Zhang, and Zhanxing Zhu for their helpful suggestions. We thank the two reviewers for helpful suggestions. This project is financially supported by Changping Laboratory and the Ministry of Science and Technology of China. X. W. was supported in part by the Peking University Boya Postdoctoral Fellowship.

